# MMR vaccination induces a trained immunity program characterized by functional and metabolic reprogramming of γδ T cells

**DOI:** 10.1101/2022.11.24.516894

**Authors:** Rutger J. Röring, Priya A. Debisarun, Javier Botey-Bataller, Tsz Kin Suen, Özlem Bulut, Gizem Kilic, Valerie A. C. M. Koeken, Andrei Sarlea, Harsh Bahrar, Helga Dijkstra, Heidi Lemmers, Katharina L. Gössling, Nadine Rüchel, Philipp N. Ostermann, Lisa Müller, Heiner Schaal, Ortwin Adams, Arndt Borkhardt, Yavuz Ariyurek, Emile J. de Meijer, Susan Kloet, Jaap ten Oever, Katarzyna Placek, Yang Li, Mihai G. Netea

## Abstract

The measles, mumps and rubella (MMR) vaccine protects against all-cause mortality in children, but the immunological mechanisms mediating these effects are poorly known. We systematically investigated whether MMR can induce long-term functional changes in innate immune cells, a process termed trained immunity, that could at least partially mediate this heterologous protection. In a randomized placebo-controlled trial, 39 healthy adults received either the MMR vaccine or a placebo. By using single-cell RNA-sequencing, we found that MMR caused transcriptomic changes in CD14-positive monocytes and NK cells, but most profoundly in γδ T cells. Surprisingly, monocyte function was not altered by MMR vaccination. In contrast, the function of γδ T cells was significantly enhanced by MMR vaccination, with higher production of TNF and IFNγ, as well as upregulation of cellular metabolic pathways. In conclusion, we describe a new trained immunity program characterized by modulation of γδ T cell function induced by MMR vaccination.

**One-sentence summary:** MMR vaccination induces cellular and metabolic reprogramming in γδ T cells towards a more active phenotype.

## INTRODUCTION

Vaccines are developed to target specific pathogens. However, an accumulating body of evidence suggests that certain live-attenuated vaccines provide a broad spectrum of protection against heterologous infections as well (non-specific beneficial effects; NSEs)(*1, 2*). Vaccine-induced NSEs are accompanied by epigenetic and metabolic changes in innate immune cells, also known as trained immunity (*3–6*). Recent studies suggest that induction of trained immunity is an attractive strategy to boost broad protection against infections and as anti-cancer therapy (*7*). Most of the studies aiming to study trained immunity induced by vaccines in humans have used the tuberculosis vaccine Bacille Calmette-Guérin (BCG) (*8–10*), while very little is known about the effects of other vaccines on innate immune cells. Although there is ample epidemiological evidence that other live attenuated vaccines also have NSEs (*2*), their potential effects on innate immune cells have not been studied.

Measles containing vaccines (MCVs) are one such group of vaccines associated with beneficial heterologous effects (*11*). Measles vaccines are routinely used in childhood immunization programs worldwide and were recently re-confirmed to be safe and effective (*12*). MMR vaccine is composed of a live-attenuated negative stranded measles virus, combined with mumps and rubella, and provides life-long immunity against measles after two doses. Several studies have confirmed higher child survival and lower morbidity after measles immunization, independent of measles-attributable disease (*13, 14*). This suggests that MMR induces trained immunity that can provide broad heterologous protection (*15, 16*). Here, we assess the potential of MMR vaccination to induce trained immunity against SARS-CoV-2 and a range of other microbial stimuli. We used single-cell multi-‘omics approaches to compare cellular heterogeneity in a randomized placebo-controlled trial of MMR re-vaccination in Dutch adults. Surprisingly, we found that MMR vaccination caused transcriptomic and functional changes in γδ T cells, rather than monocytes. This suggests that γδ T cells might have a key role in the mechanisms underlying MMR-induced trained immunity.

## RESULTS

### Study design and baseline characteristics

To investigate the potential non-specific effects of MMR re-vaccination in adults, we conducted an exploratory randomized controlled trial (**Fig. 1A**; see methods for more details). Briefly, thirty-nine healthy adults (19 female and 20 male) were randomly assigned to receive either the MMR vaccine or a placebo. All participants were between 18 and 50 years of age and there were no significant differences in sex, age, or BMI between the vaccination arms (**Fig. S1**; **Table S1**). Peripheral venous blood was collected to conduct immunological analyses was collected immediately before vaccination and one month later. No infections occurred between visits or at least two weeks prior to the baseline measurement.

**Fig. 1.**
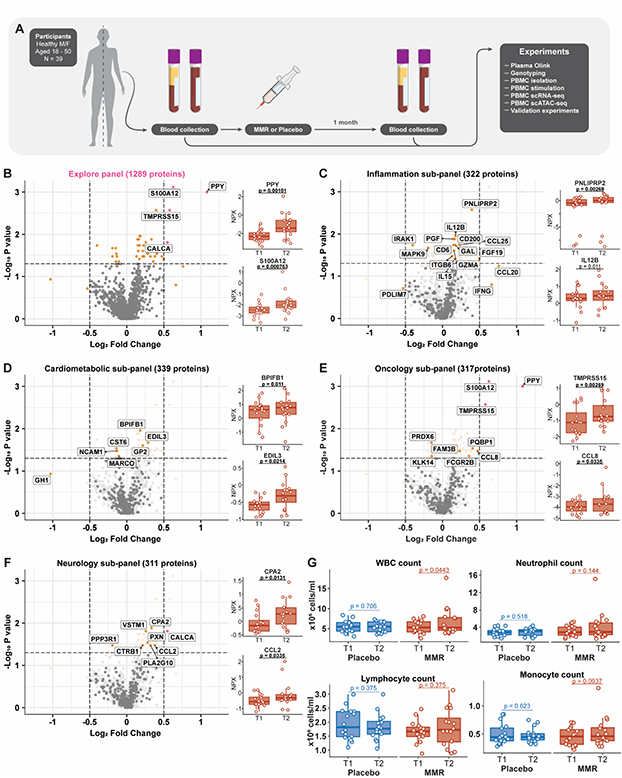
Study setup, plasma proteomics analysis after MMR vaccination, and white blood cell counts. (A) Setup of the present randomized, placebo-controlled trial of MMR vaccination. (B) Volcano plot of Olink targeted proteomics (1289 analyzed proteins in total) in plasma, after MMR vaccination (n = 16). (C-F) Volcano plots of sub-categories of the plasma proteome measured by proximity extension assay technology (Olink). In volcano plots for sub-panels, the full panel is depicted in light gray as the background. The side plots of B-F show the relative expression values (NPX) of selected proteins in each (sub)panel. (G) Total- and differentiated white blood cell counts (WBC), before and after vaccination in placebo and MMR-vaccinated groups. The p-value cutoff for the volcano plots is unadjusted p < 0.05. WBC, white blood cell; T1, baseline; T2, one month after treatment.

### Circulating biomarkers of inflammation after MMR vaccination

Previous studies have shown that trained immunity induced by BCG increases the responsiveness of innate immune cells upon rechallenge, but reduces systemic inflammation during homeostasis(*8*). We sought to investigate if this is also the case after MMR vaccination. To that end, we used Proximity Extension Assay technology (Olink) to assess targeted proteomics biomarkers (which have previously been related to inflammation, oncology, neurology, or cardiometabolic function) before and after MMR vaccination (**Fig. 1B**). There were no major changes in plasma proteome composition between the baseline measurement and 1 month after vaccination. Of all analyzed parameters (1289 after quality control), only 4 met our selected cutoffs for log2 fold change > 0.5 and unadjusted p-value < 0.05. These were PPY (a pancreatic protein associated with counter-regulation of gastric emptying(*17*)), S100A12 (a calcium-binding alarmin protein(*18*)), TMPRSS15 (a peptidase known to activate trypsin(*19*)), and CALCA (a vasodilating peptide hormone involved in calcium regulation and thought to also function as a neurotransmitter(*20*)).

We subsequently assessed the proteins that met the statistical significance threshold, independently of the fold change (**Fig. 1C-F**). Of the protein subcategories, the inflammation-related proteins showed the highest number of changes, with a trend towards upregulation after vaccination (**Fig. 1C**). PNLIPRP2, the top significant hit from this sub-panel, is a pancreas-associated protein involved in lipid metabolism. Among the other enriched proteins were factors related to NK cell and T cell activities (IL-12B, IL-15, IFNγ, CCL20, GZMA). Of the cardiometabolic-related proteins (**Fig. 1D**), most suggestive hits were also related to immunological processes either directly (GP2, CST6, NCAM1, MARCO) or indirectly (GH1). The proteins significantly changes in the oncology panel (Fig. 1E) were similarly enriched for immunologically relevant proteins (S100A12, PQBP1, CCL8, FCGR2B), with a trend towards upregulation after vaccination. Finally, in the neurology panel (**Fig. 1F**) there was also an upregulation of immunology-related proteins such as CCL2, VSTM1, and PLA2G10. These results indicate that inflammation-related proteins tend to be upregulated in the circulation after MMR vaccination, although the effect size is relatively limited. However, in terms of the effects of MMR vaccination on the systemic inflammatory status, this differs significantly from the inhibitory effects exerted by BCG vaccination.

We then considered the effects of MMR vaccination on circulating leukocyte counts (**Fig. 1G**). MMR vaccination, but not placebo treatment, significantly increased (p = 0.04) the number of circulating leukocytes after one month, although there was a large inter-individual variation. This change appeared to be driven mainly by an increase in myeloid cells (neutrophils and monocytes), although the comparison did not reach statistical significance for the individual cell types. Together, our results show that MMR-vaccinated individuals present with slightly increased systemic inflammation one month after vaccination.

### Transcriptome effects in monocytes and γδ T cells after MMR vaccination

We decided to further investigate the effects of MMR vaccination on the composition and cellular states of circulating leukocytes. To that end, we performed single-cell (sc)ATAC-seq and scRNA-seq on peripheral blood mononuclear cells (PBMCs) of a subset of participants. For scATAC-seq, we selected 12 participants from both the placebo and MMR-treated groups. We then selected half of those individuals to also perform scRNA-seq, taking care to balance male and female participants from both the MMR and placebo groups.

An integrated analysis of both scRNA-seq and scATAC-seq data (**Fig. 2A, Fig. S2A-B**) based on marker genes enabled us to identify 13 cell types within the PBMC fraction (**Fig. S2C**). Since the white blood cell differential (**Fig. 1G**) suggested there might be changes in the PBMC composition, we leveraged the single-cell datasets to investigate this in more detail. Indeed, we observed intra-individual shifts in PBMC composition between timepoints (**Fig., Fig. S3A**). However, we did not observe any consistent changes in either the MMR or placebo group (**Fig. 2B, S3A-B**). Although the low number of participants hindered statistical comparisons, our data suggest that it is unlikely that MMR vaccination has a major influence on PBMC composition one month after vaccination.

**Fig. 2.**
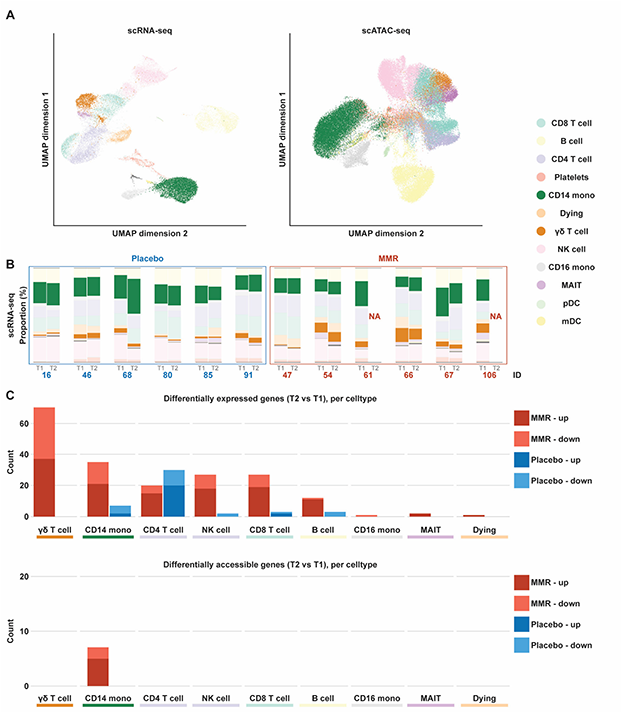
Single-cell analysis of PBMCs following MMR vaccination. (A) UMAP analysis of single-cell RNA-seq (left) and single-cell ATAC-seq of PBMCs (right). (B) Proportions of cell types annotated according to the scRNA seq data in placebo and MMR samples. (C) Differentially expressed genes (scRNA seq) per cell type, between baseline (T1) and one month after placebo or MMR (T2). (D) Differentially accessible genes (scATAC-seq) per cell type, between baseline (T1) and one month after placebo or MMR (T2).

Previous studies have shown that vaccination with other live-attenuated vaccines such as BCG induces transcriptional and functional changes (trained immunity) in innate immune cells such as monocytes (*21, 22*). We therefore hypothesized that although the PBMC composition was not altered by MMR vaccination, changes in monocyte transcriptional and functional programs could account for its known NSEs, and possibly also for our observation that MMR increases systemic inflammation. We thus assessed the transcriptional effects of MMR and placebo between timepoints, across different cell types. In every cell type except CD4^+^ T cells, MMR vaccination had a more pronounced transcriptional effect than placebo administration (**Fig. 2C**; top panel). Indeed, CD14^+^ monocytes and NK cells, classical executors of BCG-induced trained immunity, were prominently influenced by MMR. Surprisingly however, γδ T cells were the most strongly affected cells at transcriptional level among all assessed cell types. Cell-type specific analysis of differentially accessible genes revealed the chromatin to be impacted by MMR vaccination only in CD14^+^ monocytes (**Fig. 2C**; bottom panel).

### MMR has minor effects on the transcriptome and epigenome of monocyte subpopulations

Exploring further the monocyte sequencing data, we found different subpopulations defined by their transcriptional or open-chromatin signatures. Among them, we found a subpopulation highly expressing HLA genes and another one upregulating alarmins, in both data layers (**Fig. 3A**). We did not identify significant changes in the subpopulations in either data layer (**Fig. S3A**). Investigating the transcriptional changes induced by MMR, we identified more than 20 differentially expressed genes (adjusted p-value < 0.05) and 7 genes with changed chromatin accessibility (**Fig. S4C**). Upregulated genes were enriched in pathways associated with response to mechanical stimulation or exposure to metals such as calcium (*FOS*, *FOSB*, *JUN*, *JUNB* and *NFKBIA*). Downregulated genes were enriched in cell-cell adhesion programs (*B2M*, *FGL2*, *CD46*, *PRKAR1A*) (**Fig. 3B,C**). Thus, while monocytes were moderately affected at the transcriptional level by MMR vaccination, these cells were not explicitly more pro-inflammatory during homeostasis.

**Fig. 3.**
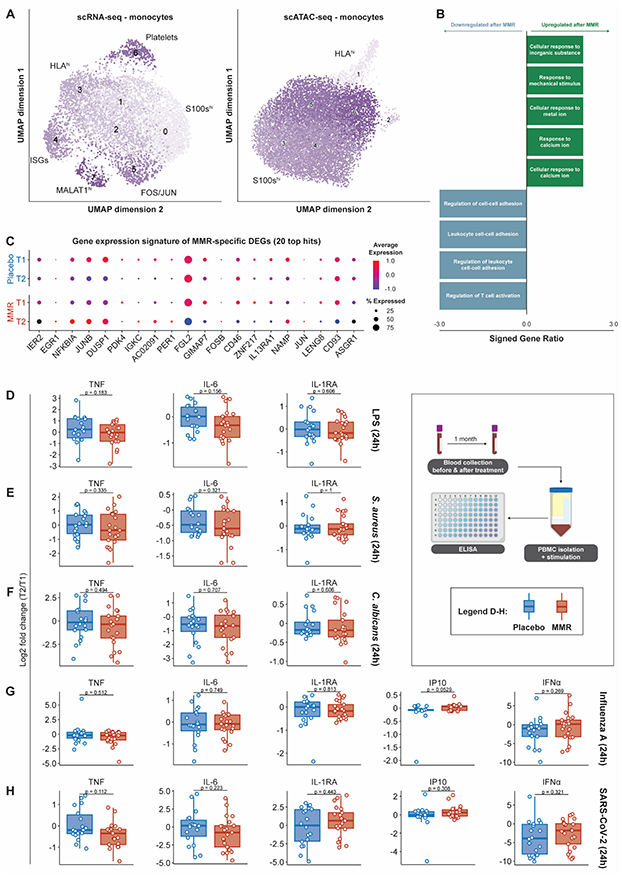
Single-cell analysis of monocyte subpopulations, and monocyte-associated cytokine production by PBMCs. (A) UMAP analysis and sub-population identification of single-cell RNA-seq (left) and single-cell ATAC-seq (right), specifically in monocytes. (B) Pathway enrichment of genes that are differentially expressed in monocytes after MMR vaccination. (C) Top 20 differentially expressed genes in monocytes following MMR vaccination, by timepoint and treatment group. (D-H) Monocyte-associated cytokines produced by PBMCs following diverse stimulations; the data are expressed as log2 fold-changes between baseline and one month after treatment.

### Cytokine production capacity of PBMCs after MMR vaccination

Innate immune memory is defined as the long-term functional reprogramming of innate immune cells by a first stimulation, leading to an altered response towards re-stimulation. To investigate functional immune responses after MMR vaccination or placebo, we stimulated PBMCs for 24 hours with a variety of bacterial (LPS, heat-killed *Staphylococcus aureus*), fungal (heat-killed *Candida albicans*), and viral (poly(I:C), R848, Influenza A (H1N1), SARS-CoV-2) stimuli, and measured monocyte-associated cytokine responses by ELISA. We measured TNF, IL-6, and IL-1RA for all stimuli. Additionally, we measured IP10 and IFNα specifically for the viral stimulations. We calculated log2-transformed fold changes corrected for age, sex, and BMI between one month after vaccination and baseline measurements, and statistically compared placebo treatment to MMR vaccination (**Fig. 3B-H**). We observed large intra- and inter-individual variations in cytokine responses between baseline and one month after treatment. However, there were no significant differences between placebo and MMR for any measured cytokine across all stimuli. Thus, we found no differences in PBMC cytokine production capacity after MMR vaccination. As monocytes are the main producers of the measured cytokines in PBMCs during 24-hour stimulations, this suggests that monocyte cytokine secretion capacity is not changed by MMR vaccination.

### Transcriptional and functional reprogramming of Vδ2 T cells after MMR vaccination

Because the most prominent changes in response to MMR were seen in γδ T cells according to the scRNA-seq data (**Fig.2**), in the next set of experiments we assessed whether vaccination changed the transcriptional and functional programs of these cells. A closer look into the γδ T cell subpopulations revealed differences in their transcriptional and open-chromatin dynamics. Transcriptionally, two distinct subpopulations were found, one characterized by transcription of granzyme genes, and the other by upregulation of IL7R (**Fig. 4A**; left). Integrating open-chromatin landscape with their transcriptional profiles from the same individuals led to differentiating Vδ1 (higher in GZMB and GZMH) and Vδ2 (higher in GZMK) T cell populations (**Fig. 4A**; right). Notably, the vaccination did not affect the proportion of the subpopulations identified (**Fig. S5A-B**). Interestingly however, MMR induced a metabolic shift in γδ T cells at the transcriptional level, with downregulation of genes involved in cellular respiration and ATP synthesis (**Fig. 4B-C**), exemplified by *NDUFA3*, *ATP5F1E*, *ATP5MD*, and *ATP5MG*. We subsequently investigated the functional consequences for Vδ2 T cells (the most abundant γδ-T cell population in the blood) by flow cytometry.

**Fig. 4.**
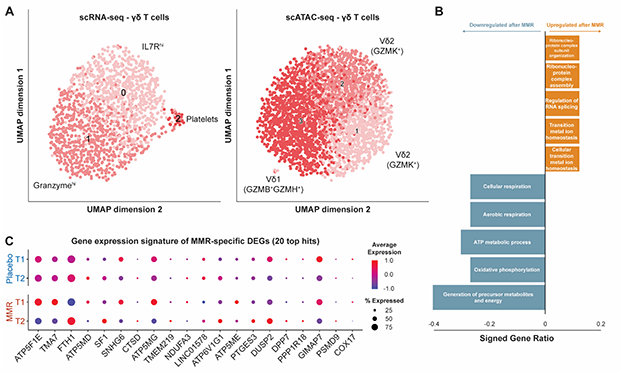
Single-cell analysis of γδ T cell populations. (A) UMAP analysis and sub-population identification of single-cell RNA-seq (left) and single-cell ATAC-seq (right), specifically in γδ T cells. (B) Pathway enrichment analysis of genes that are differentially expressed in γδ T cells after MMR vaccination. (C) Top 20 differentially expressed genes in γδ T cells following MMR vaccination, by timepoint and treatment group.

There was a small but significant decrease in Vδ2 T cells present in the PBMC fraction of MMR-vaccinated individuals (p = 0.0425); a pattern that was not present in the placebo controls (**Fig. 5A**). On the other hand, we did not find any significant differences in relative abundance of subpopulations expressing CD27 and/or CD45RA, indicating MMR did not have strong effects on classical memory-Vδ2 T cell formation (**Fig. S6A**). There were also no changes in the expressions of CTLA4, PD1, LAG3, and TIM3, markers that are commonly associated with T cell dysfunction or exhaustion. Notably, following stimulation of the γδ T cell receptor using anti-CD3/anti-CD28 beads, the percentage (but not the mean fluorescence intensity [MFI]) of Vδ2 T cells positive for TNF or IFNγ increased (**Fig. 5B, Fig. S6B**). This indicates that the γδ T cell population has become more responsive towards secondary stimulation, a feature resembling the classical monocyte trained immunity. Unstimulated Vδ2 T cells showed a trend towards lower expression of the degranulation marker CD107a after MMR vaccination (**Fig. S6C**); this effect was not present in CD3/CD28-stimulated Vδ2 T cells, which showed no difference between timepoints in the percentage of cells staining positive for CD107a, granzyme B, or perforin (**Fig. 5C**). These data suggest that granule release by Vδ2 cells is more tightly regulated at baseline following MMR vaccination, but that this effector function is not weaker following stimulation.

**Fig. 5.**
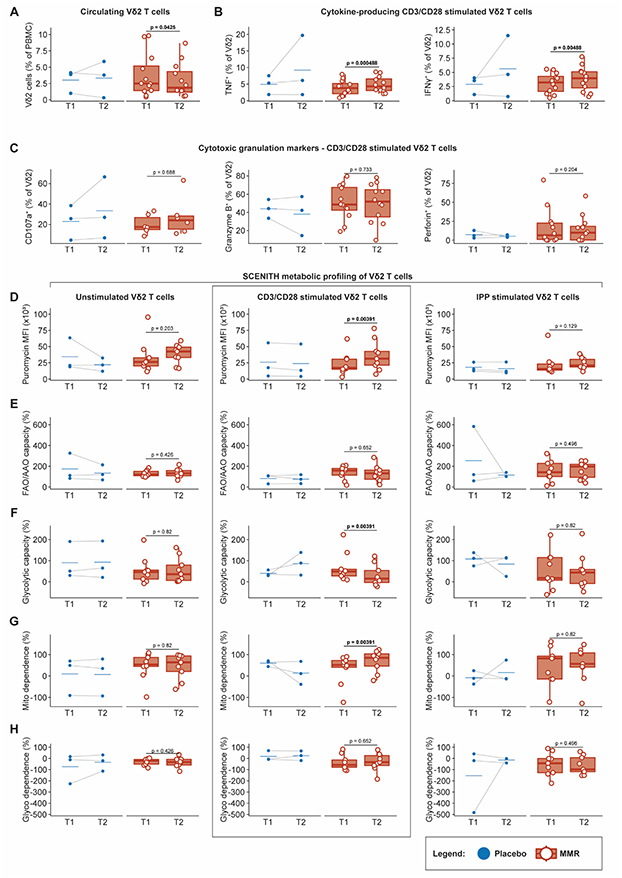
Functional and metabolic characterization of Vδ2 cells following MMR vaccination.. (A) The percentage of Vδ2 T cells in isolated PBMCs. (B) The percentage of Vδ2 T cells that produce TNF or IFNγ following CD3/CD28 stimulation. (C) The percentage of Vδ2 T cells expressing markers of cytotoxic granule release (CD107a, left) or production (Granzyme B and perforin, middle and right). metabolic parameters by modified SCENITH™ (https://www.scenith.com) calculated as in Argüello *et al*. (*23*): (D) puromycin incorporation, (E) FAO/AAO capacity, (F) Glycolytic capacity, (G) Mitochondrial dependence, (H) Glycolysis dependence. All parameters were measured by flow cytometry.

The single-cell analyses revealed that unstimulated γδ T cells modulate genes associated with oxidative phosphorylation (**Fig. 4B-C**). Therefore, we determined the functional metabolism profile of Vδ2 T cells before and after MMR vaccination using SCENITH (*23*)). This flow cytometry-based technique uses puromycin incorporation as a proxy for protein synthesis activity, which in itself reflects a significant portion of total ATP used by the cell. Briefly, we measured protein synthesis levels (puromycin MFI; **Fig. 5D**) upon treatment with metabolic inhibitors. This allowed us to calculate fatty acid/amino acid oxidation capacity (**Fig. 5E)**, glycolytic capacity (**Fig. 5F**), mitochondrial dependence (**Fig. 5G**), and glucose dependence (**Fig. 5H**) of Vδ2 T cells. These experiments revealed a trend towards higher protein synthesis activity (p = 0.203) in unstimulated Vδ2 T cells following MMR vaccination. The same pattern was visible following stimulation with isopentenyl pyrophosphate (IPP, a common antigen specifically for Vδ2 T cells; p = 0.129), and was statistically significant following CD3/CD28 stimulation (p = 0.00391). The metabolic processes fueling this change remained unclear however, as only CD3/CD28-stimulated Vδ2 T cells had changed metabolic parameters: a decrease in glycolytic capacity and thus a concomitant increased dependence on mitochondrial energy metabolism. In conclusion, these data show, for the first time, transcriptional and functional changes consistent with induction of trained immunity in γδ T cells following MMR vaccination.

## DISCUSSION

Trained immunity entails the process of boosting innate immune function following vaccination or infection (*4*), and this process has been proposed to mediate at least in part the heterologous protective effects of live attenuated vaccines such as BCG or MMR. While extensive studies have documented induction of trained immunity by BCG (including but not limited to (*3, 5, 6, 8–10, 22, 24, 25*)), nothing is known regarding the capacity of MMR vaccination to induce trained immunity. We performed a randomized, placebo-controlled trial investigating the potential of the MMR vaccine to induce innate immune memory. Using single-cell multi-omics transcriptional and epigenetic analysis combined with functional immunological and metabolic assays, we show that MMR vaccination induces a trained immunity phenotype in γδ T cells, while it has limited effects on monocyte function.

Most studies on the heterologous protection induced by certain vaccines such as BCG or influenza have focused on the induction of trained immunity in myeloid cells (*3, 26*). MMR vaccination has beneficial heterologous effects on overall mortality in children, therefore, we hypothesized that it also induces trained immunity. MMR induced some modest changes in the chromatin accessibility and transcriptional programs of monocytes, related to cellular responses to metal ions (specifically calcium; upregulated) and cellular adhesion (downregulated). However, these transcriptional effects did not result in significant effects on monocyte-derived cytokine production, despite the well-known role of calcium-dependent signaling in the function of immune cells (*27–29*). Future studies should be conducted to analyze in more depth the role of these pathways in monocytes after MMR vaccination, and if these changes are functionally relevant.

Instead, we discovered that MMR induced much stronger transcriptomic and functional changes in the innate lymphoid population of γδ T cells. In this respect, MMR vaccination modulated the expression of genes involved in aerobic energy metabolism. We sought to functionally validate these findings and therefore closely examined Vδ2 T cells, the most abundant γδ T cell subpopulation in human peripheral blood. MMR vaccination preceded a significant increase in the proportion of Vδ2 T cells producing TNF and IFNγ, and these cells were more metabolically active, especially after CD3/CD28 stimulation. Our findings indicate that γδ T cells have a more activated phenotype in MMR-vaccinated individuals, providing a plausible mechanistic explanation for the NSEs conferred by MMR.

Because of their non-canonical antigen-recognition mechanisms, the exact receptors and pathways that trigger the effects of MMR vaccination on γδ T cell transcriptome and function remain to be investigated by future studies. Previous gene expression and functional analyses demonstrate that γδ T cells have hybrid innate- and adaptive immune functions; the single-cell transcriptome of these cells has similarities to both CD8^+^ T cells and NK cells (*30*). Their innate-like features encompass the ability to mediate antibody-dependent cellular cytotoxicity, phagocytose pathogens, and direct rapid non-specific responses against threats (*31*). On the other hand, classic adaptive features include somatic recombination of their functional T cell receptor, memory cell formation, and professional antigen-presenting capabilities (*32*). Unlike classical αβ T-cell receptor signaling antigen-recognition by γδ T cells is not MHC-restricted (*33*). Vδ2 T cells, for example, predominantly recognize phosphoantigens such as isopentenyl pyrophosphate, (E)-4-Hydroxy-3-methyl-but-2-enyl pyrophosphate (HMBPP) in the context of butyrophilins 2A1 and 3A1 ((*34*)), and tetanus toxoid (*35, 36*). In cancer immunology, γδ T cells are known to exert strong anti-tumor effects by the release of pro-inflammatory cytokines, granzymes, perforin and via activation of apoptosis-triggering receptors (*36*), although pro-tumor effects have also been described ((*37*)). Interestingly, previous studies have suggested that IFNγ-producing γδ T cells are more dependent on glycolysis than on oxidative metabolism (*38*), whereas our analyses show increased reliance on mitochondrial metabolism and simultaneously an increase in IFNγ-producing Vδ2 T cells. This contrast could potentially be explained by the differences in used models: our study uses human samples rather than mice, and the previous work was done in a cancer model. Future studies need to confirm our results and assess the full array of pathways and functional consequences induced by MMR on γδ T cells.

Our data have several practical and theoretical implications. On the one hand, the differences between the BCG-induced (myeloid-dependent) and MMR-induced (lymphoid-dependent) trained immunity programs demonstrate that different vaccines can induce different types of innate immune memory. Remarkably, even when considering effects on lymphoid cells with innate properties, a major difference emerges between MMR-induced effects on γδ T cells rather than BCG effects on NK cells. On the other hand, demonstrating that MMR is also able to induce trained immunity could lead to the hypothesis that it may have beneficial heterologous effects in other groups of individuals with increased susceptibility to infections. Indeed, MMR has been proposed as a potential approach to prevent COVID19 in the period before the SARS-CoV-2 specific vaccines were available (*39, 40*). In a case-control study during a recent measles outbreak, a reduced COVID19 incidence was detected in men vaccinated with MMR (*41*).

Not only induction of trained immunity has been proposed as a mechanism explaining the heterologous effects of MMR on COVID-19. An inverse correlation between COVID-19 disease severity and MMR-specific IgG-titers was found in adults in the United States (*42*). A potential explanation for this observation could lie in cross-reactivity against structurally similar components of SARS-CoV-2 and MMR epitopes, which was described by Marakosova et al (*43, 44*). However, this cannot account for the entire breadth of protection offered by MMR vaccines and these studies did not investigate the potential impact of MMR on innate immune cells.

Our study also has some limitations. First, the sample size was limited due to the exploratory nature of this investigation, which barred us from investigating the host and environmental factors that impact these effects of MMR vaccination. Future research should encompass an increased number of participants and a broader range of study parameters such as microbiome constituents, epigenetic histone modifications, and more follow-up timepoints, similar to the large-scale vaccination studies performed with BCG (*8, 25*). Second, while NK cell activation related inflammatory proteins were increased in the serum of MMR-vaccinated individuals, we could not substantiate this finding with functional NK cell experiments due to limitations in the cell numbers available. Third, it is unknown which MMR components triggered γδ T cell activation, or if they reacted for example upon interaction with other activated cells. Moreover, in this study we in fact re-vaccinated adults who had previously received the MMR vaccine as part of the Dutch national vaccination program. Future studies should investigate if primary MMR vaccination has similar effects.

In conclusion, MMR is the first described vaccine that induces a program of trained immunity based on long-term transcriptional and functional changes of γδ T cells. The immunological and metabolic cellular responses to MMR reveal γδ T cells as a novel population of innate-like cells that mediate trained immunity. Our findings warrant further research to investigate the possibility that γδ T cells activation may be a component of trained immunity programs of other vaccines as well, and to assess the potential to improve vaccine efficacy by inducing these effects.

## Acknowledgements

We thank the volunteers of the BCG-PLUS cohort for their participation in this study. In addition, we thank our team of research nurses from the Radboud Technology Center Clinical Studies for aiding in participant visits. We are very grateful that Diletta Rosati prepared heat-killed *Candida albicans* for use in PBMC stimulation experiments. Likewise, we thank Jelle Gerretsen for preparing the heat-killed *S. aureus* we used in PBMC stimulation experiments.

## Author contributions

RJR, PAD, and JBB contributed equally to this work, and each has the right to list themselves first in author order on their CVs.

## CRediT statement

Conceptualization: MGN, YL, RJR, PAD, JBB, KP

Data Curation: PAD, RJR, JBB, OB, GK

Formal analysis: JBB, RJR, PAD, OB, GK, VACMK

Funding acquisition: YL, MGN, RJR

Investigation: PAD, RJR, TKS, OB, GK, AS, HB, HD, HL, EJdM, YA

Methodology: PAD, RJR, JBB, SK, YL, KP, MGN

Project administration: PAD, RJR, JtO, MGN

Resources: RJR, PAD, JBB, KP, EJdM, KLG, NR, PNO, LM, HS, OA, AB, YA, SK, YL, MGN

Software: JBB, RJR, PAD, OB, VACMK, YL

Supervision: YL, MGN

Verification: RJR, JBB, TKS, KP

Visualization: RJR, JBB

Writing – original draft: RJR, PAD, JBB, MGN

Writing – review & editing: RJR, PAD, JBB, MGN

## Data and materials availability

The sequencing data used in this manuscript will be made accessible in the EGA archive (EGAS00001006787). Olink data will be added as a supplementary file. Other data are available upon request to the corresponding author.

## Supplementary materials

Materials and methods

Figs. S1 – S6

Tables S1 – S3

References: 45-55

## Supplementary materials

### MATERIALS AND METHODS

#### Study design

This randomized placebo-controlled trial, depicted in Fig. 1A, was designed to research the ability of MMR vaccination to establish trained immunity. Therefore, participants were 1:1 allocated to receive either a placebo vaccination (0.1 ml of 0.9% saline solution), or an MMR vaccination (SD, 0.5 ml, live attenuated mumps virus [strain ‘Jeryl Lynn’, at least 12.5 * 10^3 CCID50]; live attenuated measles virus [strain ‘Enders’ Edmonston’, at least 1 * 10^3 CCID50]; live attenuated rubella virus [strain ‘Wistar RA 27/3’, at least 1 * 10^3 CCID50]). Vaccination was performed intramuscularly in the right upper arm and. Blood was drawn at baseline (“T1”) and one month after vaccination (“T2”). The trial protocol registered under NL74082.091.20 in the Dutch trial registry, was approved in 2020 by the Arnhem-Nijmegen Ethics Committee. All experiments were conducted in accordance with the Declaration of Helsinki and no adverse events were recorded.

#### Study subjects

Thirty-nine healthy volunteers (**Table S1**) within the age of 18 and 50 years, were recruited between June and September 2020. Subjects with a medical history associated with immunodeficiency or a solid or non-solid malignancy within the two preceding years were excluded. Vaccination three months prior to the start of the study or plans to receive other vaccinations during the study period was not allowed. Acute illness within two weeks before study initiation or the use of drugs, including non-steroidal anti-inflammatory drugs (NSAIDs) less than four weeks before the start of the trial, with the exception of oral contraceptives, also resulted in exclusion. Pregnant subjects were not eligible. All participants gave written informed consent.

#### Blood collection and sample processing

EDTA whole blood (8 × 10 ml) was collected via venipuncture. Two of the EDTA tubes were centrifuged immediately after collection at 2970 x *g* for 10 minutes at room temperature (RT) and plasma was stored at −80 °C until later analysis. Hematological parameters such as white blood cell count and differential were measured on a Sysmex XN-450 apparatus. Additionally, 1 ml of whole blood was stored at −80 °C for genotyping analysis.

Peripheral blood mononuclear cells (PBMCs) were isolated by density-gradient centrifugation over Ficoll-Paque. Briefly: EDTA blood was diluted in calcium/magnesium-free PBS and layered on Ficoll-Paque solution. After centrifugation for 30 minutes at 615 x *g* (no brakes; RT), the PBMC layer was collected and washed at least 3 times with cold calcium/magnesium-free PBS. The cells were resuspended in RPMI-1640 with Dutch modifications (Invitrogen) supplemented with 50 mg/ml gentamicin (Centrafarm), 2 mM GlutaMAX (Gibco) and 1 mM pyruvate (Gibco) and counted by Sysmex. For cytokine production assessments, PBMCs were seeded in round-bottom 96 well plates at 0.5 * 10^6^ cells/well. The cells were stimulated for 24 hours using the stimuli described in **Table S3** (all in the presence of 10% human pooled serum, at 37 °C and 5% CO_2_). Supernatants were collected and stored at −20 °C until further analysis.

#### PBMC freezing and thawing

Leftover PBMCs were resuspended in ice-cold, heat-inactivated fetal bovine serum (FBS) prior to cryopreservation. Ice-cold 20% DMSO in FBS was added dropwise to the cells until a final concentration of 10% DMSO was reached. The cells were stored for up to 24 hours in CoolCell alcohol-free freezing containers (Corning) at −80 °C, after which they were transferred to a −150 °C freezer for long-term storage.

For subsequent experiments, PBMCs were thawed following a protocol modified from Hønge et al (*45*). The PBMCs were retrieved from the −150 °C storage and kept on dry ice until the moment of thawing. The cells were rapidly warmed in a water bath of 37 °C until only a small clump of ice was present in the vial. The contents were immediately transferred into a 10x volume of pre-warmed thawing medium (RPMI supplemented as described above, further supplemented with 20% FBS and 12.5 μg/ml DNase-I). The cells were centrifuged at 500 x g for 10 minutes at RT and resuspended in thawing medium without DNase-I. The cells were again centrifuged, resuspended in cold PBS and counted with trypan blue to assess recovery and viability.

#### ELISA cytokine measurements and data analysis

The cytokines TNF (commonly referred to as TNF-α), IL-6, IL-1Ra, and IP-10 were measured using DuoSet ELISA kits from R&D systems, and IFNα with a kit from PBL Assay Science, using the manufacturer’s protocol. To account for plate-to-plate variation, the participants were randomized over different plates (timepoints were kept together on the same plate). Cytokine concentrations were calculated relative to the standard curve in Gen5 software (BioTek). Log2-fold changes were calculated between T2 and T1, and corrected for sex, age, and BMI using a linear regression approach. The MMR and placebo groups were compared using Mann-Whitney U tests.

#### Targeted proteomics analysis by proximity extension assay

Plasma samples from 16 MMR-vaccinated individuals were sent to Olink (Sweden) for targeted proteomics analysis using proximity extension assay technology. In total, 1472 proteins were measured, of which 183 were removed from the analysis due to them being poorly detectable in >25% of samples (30 cardiometabolic proteins, 46 inflammatory proteins, 56 neurology proteins, 51 oncology proteins). Unadjusted p-values were calculated using the Wilcoxon signed rank test.

#### DNA isolation and genotyping

Whole blood samples were shipped on dry ice to the Human Genomics Facility of the Genetic Laboratory of the Department of Internal Medicine at Erasmus MC, Rotterdam, The Netherlands. There, DNA isolation was performed and samples were genotyped using Illumina GSA Arrays “Infinium iSelect 24×1 HTS Custom Beadchip Kit”.

#### Single cell library preparation and sequencing

Cryopreserved PBMCs were thawed as described above and washed an additional time with ice-cold PBS. Single cell gene expression libraries were generated on the 10x Genomics Chromium platform using the Chromium Next GEM Single Cell 3’ Library & Gel Bead Kit v3.1 and Chromium Next GEM Chip G Single Cell Kit (10x Genomics) according to the manufacturer’s protocol. Single-cell ATAC-seq libraries were generated on the 10x Genomics Chromium platform using the Chromium Next GEM Single Cell ATAC Library & Gel Bead Kit v1.1 and Chromium Next GEM Chip H Single Cell Kit (10x Genomics) according to the manufacturer’s protocol. Gene expression and ATAC-seq libraries were sequenced on a NovaSeq 6000 S4 flow cell using v1.5 chemistry (Illumina).

#### Single-cell sequencing data analysis

##### Pre-processing and demultiplexing scRNA-seq and snATAC-seq data

The proprietary 10x Genomics CellRanger pipeline (v4.0.0) was used with default parameters. CellRanger count or CellRanger-atac count was used to align read data to the reference genome provided by 10x Genomics. refdata-cellranger-arc-GRCh38-2020-A-2.0.0 was used for the snATAC-seq experiments and refdata-gex-GRCh38-2020-A for the scRNA-seq. In scRNA-seq, a digital gene expression matrix was generated to record the number of UMIs for each gene in each cell. In snATAC-seq, fragment files were created.

Each library was further demultiplexed by assigning cell barcodes to their donor. Souporcell (v1.3 gb) (*46*) was used for genotype-free demultiplexing by calling candidate variants on the pre-mapped bam files. Cells were clustered by their allelic information and each cluster was matched to a donor with a known genotype.

##### scRNA-seq data analysis

The expression matrix from each library was loaded to R/Seurat package (v3.2.2) (*47*) for downstream analysis. To control the data quality, we first excluded cells with ambiguous assignments from Souporcell demultiplex. Next, we further excluded low-quality cells with > 25% mitochondrial reads, < 100 or > 3,000 expressed genes, or > 5000 UMI counts.

After QC, we applied LogNormalization (Seurat function) and scaled the data, regressing for total UMI counts, number of features, percentage of mitochondrial genes and percentage of ribosomal genes. We then performed principal component analysis (PCA) based on the 2,000 more highly variable features identified using the vst method implemented in Seurat. As batches showed a good integration of the data, no integration algorithm was applied. Cells were then clustered using the Louvain algorithm with a resolution of 0.75 based on neighbors calculated the first 30 principal components. For visualization, we applied UMAP based on the first 30 principal components.

##### Annotation of scRNA-seq clusters

Clusters were annotated by manually checking the expression of known marker genes. Cluster 5 showed a higher expression of gamma and delta chain genes (*TRGC1*, *TRDC*), along with T cell markers (*CD3E*). Performing dimensionality reduction and clustering on this subset revealed two mixed populations: MAIT cells overexpressing *KLRB1* and γδ T cells over expressing *TRDC*/*TRGC*, these cells were then annotated accordingly.

##### Differential gene expression and gene-set enrichment

For paired comparison between timepoints in both MMR and placebo, differential expression (DE) tests were performed using the FindMarkers functions in Seurat with MAST (*48*). Patient ids were regressed out, in order to perform a paired analysis. Genes with a Bonferroni-corrected P-value < 0.05 were regarded as differentially expressed.

Gene-set enrichment was performed using the enrichGO function from the R package clusterProfiler. Gene-sets enriched with a Benjamin-Hochberg corrected p-value below 0.05 and more than 4 genes were considered significant.

##### snATAC-seq data analysis

ArchR (*49*) was used for the downstream analyses on snATAC-seq data, reading the fragment files created in by CellRanger-atac. Cells with fewer than 1,000 unique fragments, a transcription start site enrichment below 4, identified as doublets by ArchR, or ambiguously labelled by souporcell were removed.

After QC, we used the ArchR function ‘addIterativeLSI’ to process iterative latent semantic indexing using the top 25,000 variable features and top 30 dimensions. For visualization, we applied UMAP with nNeighbors = 30 and minDist = 0.5.

Gene scores were calculated for each cell based on accessibility. In order to aid the analysis of gamma-delta T cells, a modified reference was used. Adding the gtf gene reference used by CellRanger, gene scores could be calculated for *TRDC*, *TRGC1* and *TRGC2*.

##### snATAC-seq annotation and integration with scRNA-seq data

ArchR function ‘addGeneIntegrationMatrix’ was used to compare the calculated snATAC-seq gene score matrix and the measured gene expression in scRNA-seqdata. This resulted in a matched scRNA profile and predicted cell type per sequenced cell in the snATAC-seq data. Cell types were therefore assigned to the snATAC-seq data based on the predicted cell type of the integration. UMAPs of the integrated blocks were inspected in order to examine the quality of the integration (**Fig. S2A-B**).

##### Per cell type analysis of snATAC-seq data

A common approach was followed to inspect the open-chromatin changes in each cell type. The same method as described before for the whole cell pool was used for visualization and clustering separately in each cell type. After, open-chromatin peaks were calculated by running ‘addReproduciblePeakSet’ using Macs2 algorithm (*50*). Transcription factor motif deviations were calculated based on the ‘CIS-BP’ database annotation (*51*) using the ‘addDeviationsMatrix’ function. The effects of MMR vaccination and placebo were assessed by running ‘getMarkerFeatures’ comparing both timepoints for all data types: open-chromatin peaks, TF motifs and gene score. FDR < 0.05 was indicative of significant changes.

#### Flow cytometry measurements of γδ T cell parameters

The flow cytometry staining was performed as follows: 5×10^5^ thawed PBMCs were stained for surface markers using the antibodies described in **Table S3**, for 30 minutes in the dark at 4 °C, in FACS buffer (PBS, 5% FBS, 2 mM EDTA Intracellular proteins were analyzed after fixation and permeabilisation in Cytofix permeabilization/fixation reagent (BD biosciences) for 30 mins. Following two washes with Cytofix permeabilization/washing buffer (BD biosciences), the cells were stained with the antibodies against intracellular markers detailed in **Table S3**, for 30 minutes in the dark at 4 °C. After completion of the staining procedure, the cells were washed with PBS and stored in CellFIX reagent (BD biosciences) until acquisition on a LSR II cytometer (BD biosciences).

The flow cytometry data was analyzed in FlowJo (vX.07). The gating strategy was as follows: events corresponding to lymphocyte size were selected based on FSC-A/SSC-A, followed by selection of single-cell events in subsequent FSC-H/FSC-A and FSC-W/FSC-A gates. Viable cells were selected by gating on viability-dye-negative cells. The analyses were performed on CD45^+^CD3^+^Vδ2^+^ cells.

For measurement of surface markers on unstimulated Vδ2 T cells, the thawed PBMC were stained as described above immediately after thawing. For measurements of cytokine expression or surface markers after stimulation, the PBMCs were first treated with soluble anti-CD3/anti-C28 (BD bioscience) for 4 hours in the presence of a Golgi plug (Brefeldin A; BD bioscience), under standard cell culture conditions.

#### SCENITH

We modified the original SCENITH™ technique (https://www.scenith.com) to analyze energy metabolism of γδ T cells. Briefly, PBMCs were plated at 0.3×10^6^ cells/well in 96-well plates. The cells were cultured in RPMI alone or stimulated with soluble anti-CD3/anti-CD28, or IPP for 4 hours under standard cell culture conditions. Cells were then untreated (control) or treated with 2-DG (final concentration 100 mM), Oligomycin (O, final concentration 10 μM) and combination of 2-DG and Oligomycin (DGO, final concentration 100 mM and 10 μM) for 30 min under standard cell culture conditions. Following addition of puromycin (final concentration 10 μg/ml), the cells were incubated for an additional 45 minutes. The cells were subsequently harvested and washed in cold FACS buffer before being stained as described above.

#### Statistical analysis and software

All data was analyzed in R as described in each relevant section of the methods. Unless otherwise indicated, two-tailed p-values of <0.05 were considered statistically significant. If correction for multiple testing was applied, the method is described in the relevant methods section. In cases where the p-value is not provided, an asterisk (*) indicates statistical significance.

The following R packages were used for the present work: the Tidyverse core packages 1.3.2 (*52*), Seurat 4.1.1, ArchR 1.0.2, SeuratObject 4.1.0, GenomicRanges 1.48.0, data.table 1.14.2, ggplot2 3.3.6, colortools 0.1.6, clusterProfiler 4.4.4. magrittr 2.0.3, ggprism 1.0.3, ggsci 2.9, rstatix 0.7.0, pzfx 0.3.0, janitor 2.1.0, readr 2.1.3, openxlsx 4.2.5, psych 2.2.9. The figures were compiled in Adobe Illustrator.

### Supplementary figures

**Figure S1:**
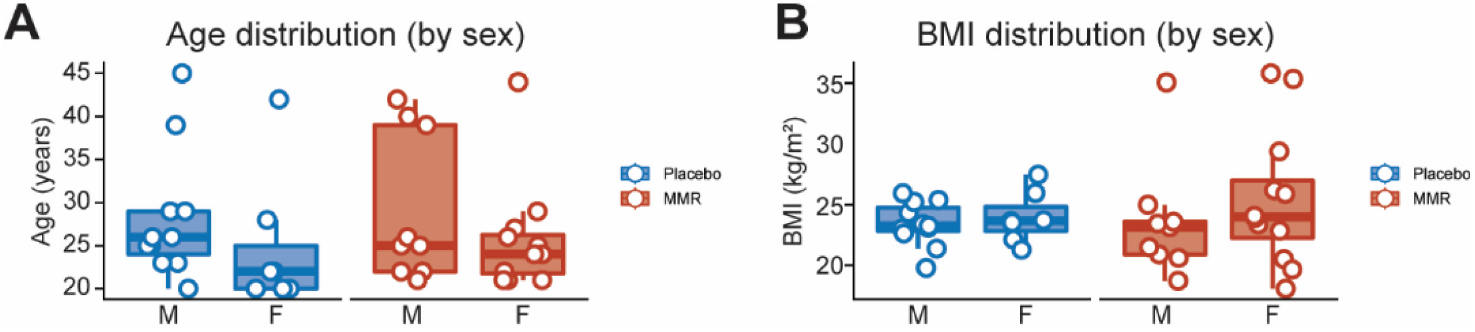
Participant characteristics. (**A**) Participant age, stratified by sex and treatment group. (**B**) Participant BMI, stratified by sex and treatment group.

**Figure S2:**
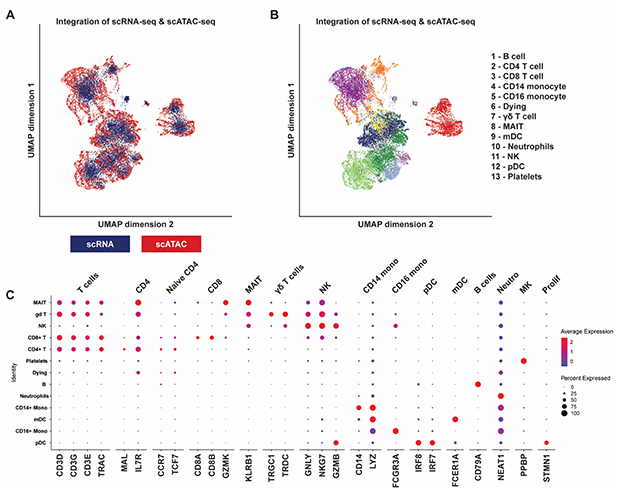
Integration of scRNA and scATAC-sequencing data and cell-type annotation. (**A**) UMAP of scRNA-seq and scATACseq integrated data. Integration was performed using canonical correlation analysis between gene expression values and gene score values calculated based on gene accessibility. () Same as before, colored by the different celltypes present. (**C**) Dotplot of markers used for celltype annotation.

**Figure S3:**
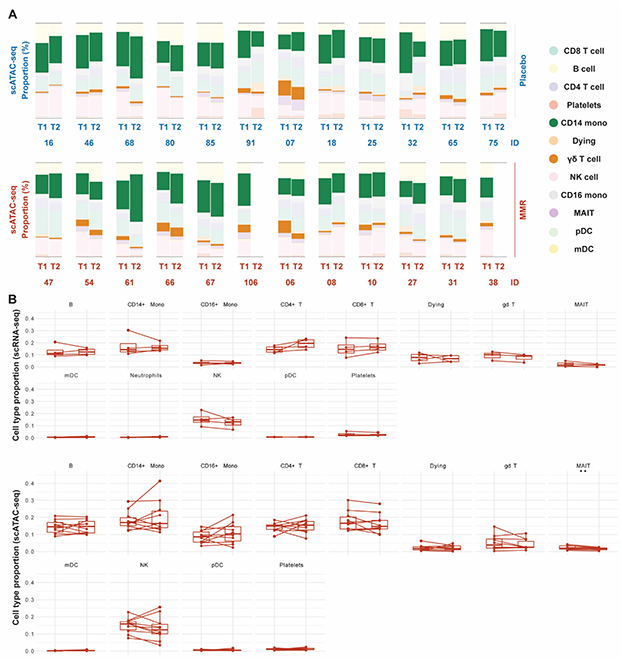
Cell-type proportions before and after treatment. (**A**) Proportions of cell types annotated according to the scATAC-seq data in placebo and MMR samples. (**B**) The same data, only displayed as boxplots per cell type to enable paired comparisons.

**Figure S4:**
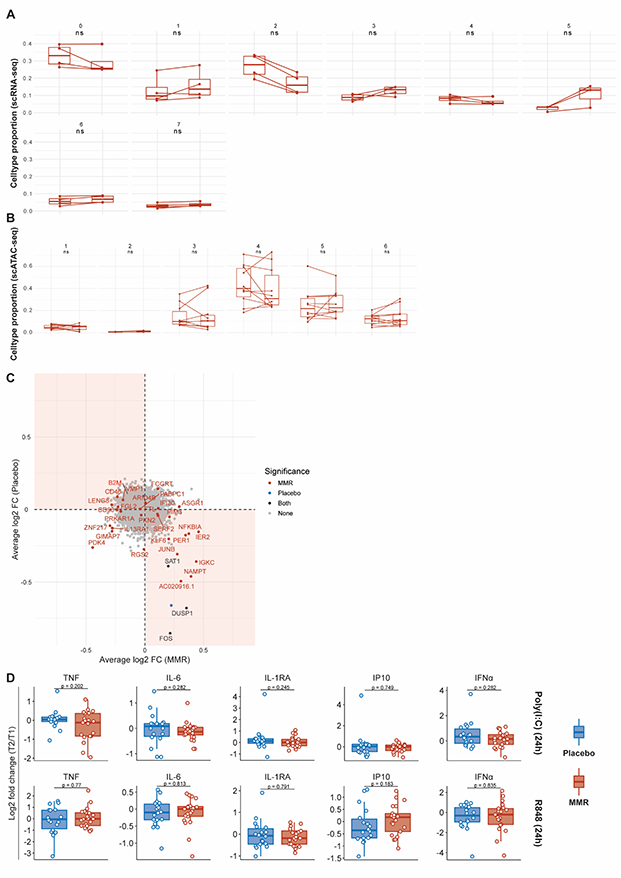
Single-cell analysis of monocyte subpopulations and monocyte-associated cytokine production by PBMCs. (**A**) Proportions of monocyte sub-populations (scRNA-seq) before and after MMR vaccination. (**B**) Proportions of monocyte sub-populations (scATAC-seq) before and after MMR vaccination. (**C**) Combined ‘volcano plot’ showing average log2 fold changes of monocyte gene expression between timepoints for both placebo and MMR. Bonferroni adjusted p-value < 0.05, paired test using MAST. (**D**) Monocyte-associated cytokines produced by PBMCs following diverse stimulations; the data are expressed as log2 fold-changes between baseline and one month after treatment.

**Figure S5:**
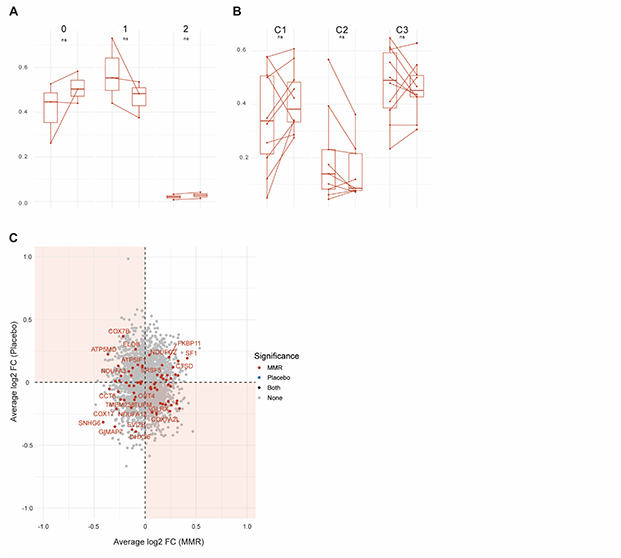
Single-cell analysis of γδ T cell populations. (**A**) Proportions of γδ T cell sub-populations (scRNA-seq) before and after MMR vaccination. (**B**) Proportions of γδ T cell sub-populations (scATAC-seq) before and after MMR vaccination. (**C**) Combined ‘volcano plot’ showing average log2 fold changes of γδ T cell gene expression between timepoints for both placebo and MMR. Bonferroni adjusted p-value < 0.05, paired test using MAST.

**Figure S6:**
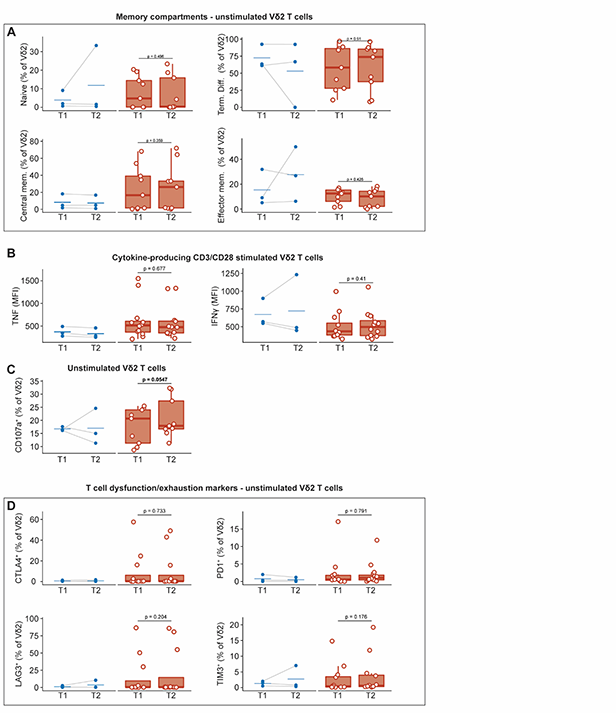
characterization of Vδ2 cells following MMR vaccination. (**A**) Proportions of memory compartments characterized by expression of CD27 and CD45RA. (**B**) Mean fluorescence intensities (MFI) of TNF and IFNγ in stimulated Vδ2 T cells. (**C**) Percentage of unstimulated Vδ2 T cells that stain positive for CD107a, a marker of cytotoxic degranulation. (**D**) Proportions of Vδ2 T cells that stain positive for markers commonly associated with T cell dysfunction (CTLA4, PD1, TIM3, LAG3).

### Supplementary tables

**Table S1:** characteristics of the study population.

| Characteristic * | Placebo<br>n = 18 | MMR<br>n = 21 | p-value† |
| --- | --- | --- | --- |
| Sex, female | 7 (38.89) | 12 (57.14) | 0.3406 |
| Sex, male | 11 (61.11) | 9 (42.86) |  |
| Age, years | 25 ± 7.63 | 25 ± 7.38 | 0.876 |
| BMI, kg/m² | 23.47 ± 1.89 | 23.45 ± 5.22 | 0.945 |
| * Median +/- SD for continuous variables and n (%) for categorical variables |  |  |  |
| † Mann-Whitney U test for continuous variables and Fisher's exact test for categorical variables |  |  |  |

**Table S2:** Stimuli used to assess PBMC cytokine production capacity.

| Stimulus | Concentration<br>in experiment | Cat. No. | Manufacturer |
| --- | --- | --- | --- |
| *E. coli* LPS serotype O55:B5,<br>further purified as in (53) | 10 ng/ml | - | Prepared in-house by<br>Heidi Lemmers,<br>Radboudumc |
| Heat-killed *S. aureus*<br>ATCC 25923 | 1 * 10<sup>6</sup> CFU/ml | - | Cultured and heat-killed in-house by Jelle<br>Gerretsen,<br>Radboudumc |
| Heat-killed *C. albicans* yeast,<br>UC820 (ATCC MYA-3573),<br>prepared as described in (54) | 1 * 10<sup>6</sup> CFU/ml | - | Prepared in-house by<br>Diletta Rosati,<br>Radboudumc |
| Poly(I:C) | 10 µg/ml | tlrl-pic | Invivogen |
| R848 | 3 µg/ml | tlrl-r848 | Invivogen |
| Influenza A H1N1, prepared<br>according to the methods in<br>(55) | 3.3 * 10<sup>5</sup> /ml | - | Kindly prepared by<br>KLG, NR, PNO, LM, HS,<br>OA, AB |
| SARS-CoV-2, prepared<br>according to the methods in<br>(55) | 1.4 * 10<sup>3</sup> /ml | - |  |

**Table S3:** Antibodies used in the flow cytometric analyses of Vδ2 T cells.

| Target | Fluorophore | Clone | Cat. No. | Manufacturer | Panel |
| --- | --- | --- | --- | --- | --- |
| Viability | Live-or-Dye Fixable Viability Stain | - | 32006 | Biotium | 1 |
| CD3 | Pacific blue | UCHT1 | 300431 | Biolegend | 1 |
| CD4 | PE-Cy7 | OKT4 | 317414 | Biolegend | 1 |
| CD8 | APC-Cy7 | SK1 | 344746 | Biolegend | 1 |
| CD45 | BV605 | HI30 | 304042 | Biolegend | 1 |
| Vδ2 TCR | FITC | REA771 | 130-111-009 | Miltenyi | 1 |
| Vδ1 TCR | PE | REA173 | 130-120-440 | Miltenyi | 1 |
| TNF | APC | MAb11 | 502912 | Biolegend | 1 |
| IFNγ | PerCP-Cy5.5 | B27 | 560704 | BD Pharmingen | 1 |
| Target | Fluorophore | Clone | Cat. No. | Manufacturer | Panel |
| Viability | Live/dead Fixable Aqua dead cell stain | - | L34957 | Invitrogen | 2 |
| CD3 | Pacific blue | UCHT1 | 300431 | Biolegend | 2 |
| Vδ2 TCR | FITC | REA771 | 130-111-009 | Miltenyi | 2 |
| Vδ1 TCR | PE | REA173 | 130-120-440 | Miltenyi | 2 |
| CD4 | BV605 | OKT4 | 317438 | Biolegend | 2 |
| CD8 | APC-Cy7 | SK1 | 344746 | Biolegend | 2 |
| PD1 | PerCP-Cy5.5 | A17188 B | 621613 | BD Bioscience | 2 |
| Perforin | Pe-Cy7 | dG9 | 308126 | Biolegend | 2 |
| Granzyme B | AF647 | GB11 | 515406 | Biolegend | 2 |
| CD45 | PE/Dazzle594 | HI30 | 304052 | Biolegend | 2 |
| Target | Fluorophore | Clone | Cat. No. | Manufacturer | Panel |
| Viability | Live/dead Fixable Aqua dead cell stain | - | L34957 | Invitrogen | 3 |
| CD3 | APC | UCHT1 | 300458 | Biolegend | 3 |
| CD4 | PE-Cy7 | OKT4 | 317414 | Biolegend | 3 |
| CD8 | APC-Cy7 | SK1 | 344746 | Biolegend | 3 |
| Vδ1 TCR | Pacific blue | REA173 | 130-100-555 | Miltenyi | 3 |
| Vδ2 TCR | FITC | REA771 | 130-111-009 | Miltenyi | 3 |
| CD107a | PE | H4A3 | 328607 | Biolegend | 3 |
| CTLA4 | BV605 | BNI3 | 369610 | Biolegend | 3 |
| Lag3 | PE/Dazzle594 | 11C3C6 5 | 369331 | Biolegend | 3 |
| Target | Fluorophore | Clone | Cat. No. | Manufacturer | Panel |
| Viability | Live/dead Fixable Aqua dead cell stain | - | L34957 | Invitrogen | 4 |
| CD45 | PE/Dazzle594 | HI30 | 304052 | Biolegend | 4 |
| CD45RA | APC | HI100 | 304111 | Biolegend | 4 |
| CD3 | Pacific blue | UCHT1 | 300458 | Biolegend | 4 |
| CD4 | PE-Cy7 | OKT4 | 317414 | Biolegend | 4 |
| CD8 | APC-Cy7 | SK1 | 344746 | Biolegend | 4 |
| Vδ1 TCR | PE | REA173 | 130-120-440 | Miltenyi | 4 |
| Vδ2 TCR | FITC | REA771 | 130-111-009 | Miltenyi | 4 |
| CD27 | PerCP-Cy5.5 | M-T271 | 560612 | BD Bioscience | 4 |
| NKG2D | BV605 | 1D11 | 320832 | Biolegend | 4 |
| Viability | Live-or-Dye Fixable Viability Stain | - | 32006 | Biotium | 5 |
| CD3 | Pacific blue | UCHT1 | 300431 | Biolegend | 5 |
| CD4 | PE-Cy7 | OKT4 | 317414 | Biolegend | 5 |
| CD8 | APC-Cy7 | SK1 | 344746 | Biolegend | 5 |
| CD45 | BV605 | HI30 | 304042 | Biolegend | 5 |
| Vδ2 TCR | APC | REA771 | 130-111-009 | Miltenyi | 5 |
| Vδ1 TCR | PE | REA173 | 130-120-440 | Miltenyi | 5 |
| Puromycin | AF488 | 12D10 | MABE343 | Merck | 5 |

